# Tactile stimulations reduce or promote the segregation of auditory streams: psychophysics and modelling

**DOI:** 10.1101/2024.12.05.627120

**Authors:** Farzaneh Darki, James Rankin, Piotr Słowiński

## Abstract

Auditory stream segregation plays a crucial role in understanding the auditory scene. This study investigates the role of tactile stimulation in auditory stream segregation through psychophysics experiments and a computational model of audio-tactile interactions. We examine how tactile pulses, synchronized with specific tones in a sequence of interleaved high- and low-frequency tones (ABA-triplets), influence the likelihood of perceiving integrated or segregated auditory streams. Our findings reveal that tactile pulses synchronized with specific tones enhance perceptual segregation, while pulses synchronized with both tones promote integration. Based on these findings, we developed a dynamical model that captures interactions between auditory and tactile neural circuits, including recurrent excitation, mutual inhibition, adaptation, and noise. The proposed model shows excellent agreement with the experiment. Model predictions are validated through psychophysics experiments. In the model, we assume that selective tactile stimulation dynamically modulates the tonotopic organization within the auditory cortex. This modulation facilitates segregation by reinforcing specific tonotopic responses through single-tone synchronization while smoothing neural activity patterns with dual-tone alignment to promote integration. The model offers a robust computational framework for exploring cross-modal effects on stream segregation and predicts neural behaviour under varying tactile conditions. Our findings imply that cross-modal synchronization, with carefully timed tactile cues, could improve auditory perception with potential applications in auditory assistive technologies aimed at enhancing speech recognition in noisy settings.

## Introduction

The difficulty in following a single voice or conversation in noisy situations is one of the main frustrations of people with hearing impairment, even when fitted with modern hearing aids (1). This difficulty arises from the challenge of separating a single sound stream from many competing sounds, a process known as auditory stream segregation (2; 3). Auditory stream segregation has been widely studied with the auditory streaming paradigm, an idealized stimulus for which two sequences of pure tones can be segregated into different streams (4; 5). Stream segregation has been shown to be influenced by cross-modal interaction (6). Visual signals have been shown to improve stream segregation ability (7). Tactile cues have also been shown to improve the ability to segregate interleaved melodies, even under challenging listening conditions (8). Tactile stimuli have the capacity to improve speech recognition in noise (9; 10; 11). However, our understanding of the underlying neural dynamics of this process is still limited.

Sensory substitution theory provides a compelling framework for understanding how one sensory modality can compensate for a deficit in one or augment another during complex perceptual tasks (12; 13). Auditory perception, for example, can be influenced by stimuli from other sensory modalities, such as vision (14). Although the impact of visual input on auditory perception has been extensively studied, the influence of somatosensory (tactile) stimuli remains less well understood (15; 16; 17). While research suggests that tactile inputs can modulate auditory perception (18; 19), it is unclear which attributes of tactile and auditory signals (e.g., frequency, intensity, rhythm, timing, duration) play the most critical role in enhancing or suppressing the combined sensory experience.

In (20) the influence of tactile distractors on the ability to discriminate the frequency and intensity of auditory tones was studied through a series of psychophysical experiments. The authors demonstrated that auditory frequency perception was systematically biased by tactile distractors: distractors at frequencies lower than that of the auditory tones induced larger bias effects than distractors at higher frequencies. They also observed that tactile distractors biased the intensity of auditory perception. The magnitude of this effect scaled with distractor intensity, but did not vary with distractor frequency (20). These findings highlight the complex interplay between tactile and auditory modalities, raising questions about the nature of cross-modal interactions between touch and sound, as well as the specific neural mechanisms underlying audio-tactile interactions.

Multisensory integration is traditionally thought to occur in higher association cortices, from which multisensory signals are relayed to (subcortical) areas involved in planning and executing actions (21; 22). According to this view, multisensory integration occurs only after unisensory information has been thoroughly processed along its specific sensory hierarchy. However, recent results challenge this notion and suggest that multisensory interactions can occur in early sensory areas. In particular, fMRI (23; 24; 25; 26; 27) and electrophysiological studies in monkeys (28; 29; 30) found multisensory activations in brain areas considered unisensory, supporting the idea that multisensory integration can occur early in sensory processing through feedforward mechanisms, independent of attention (preattentive bottom-up mechanisms (23)).

The audiotactile cross-modal interaction could occur due to both direct and indirect pathways connecting auditory and somatosensory processing areas (31). Some neurons within the primary auditory cortex (A1) respond to tactile stimuli, directly reflecting tactile processing in auditory regions (28; 32). Tactile pulses can influence tonotopic responses in the auditory cortex (33). Although A1 is traditionally associated with processing sound frequencies, studies show that it can also respond to multisensory inputs (19; 34), such as tactile, particularly when those stimuli are temporally or spatially relevant to auditory processing. Tactile stimulation, especially rhythmic pulses, can influence the timing and phase of neural oscillations in A1 (35; 36), potentially enhancing the synchronization of responses. This is more likely if the tactile input is closely synchronized with auditory input, as it can alter the salience of specific frequencies (37; 38).

Mathematical modelling has advanced our understanding of bistable phenomena including auditory streaming (39; 40) and tactile rivalry (41). A variety of models have been proposed to account for the segregation of two streams of sounds (39; 42). Rankin et al. (43; 44; 45) proposed a neuromechanistic model that captures the complex dynamics of perceptual alternations, including the dependence of perceptual durations on parameters such as frequency differences and presentation rate. The model reproduces characteristic features, such as the log-normal distribution of perceptual durations and the build-up effect. The model replicates the periodic, pulsatile responses of A1 and its dependence on stimulus features, which are pooled as inputs to a downstream competition stage. The competition between units arises from a combination of mutual inhibition, adaptation, and additive noise mechanisms, which are thought to contribute to perceptual bistability at cortical stages (46; 47; 48).

Although multisensory integration has received significant attention within the neuroscience community, the number of mechanistic mathematical models of multisensory integration remains limited. To bridge this gap, we propose a novel computational model for audio-tactile integration that can be generalized to other cross-sensory interactions. By integrating neuromechanistic modelling with psychophysics experiments, we elucidate how tactile sensations influence the perception of multiple sound sources and the underlying neural computations driving audio-tactile interactions. The proposed model extends the mathematical framework of auditory streaming. Specifically, we use the hypothesis that tactile inputs enhance excitation in the tonotopic response within the primary auditory cortex to extend the model of interactions between primary (A1) and non-primary auditory cortices presented in (43). We experimentally validate the model. Our model aligns with preattentive bottom-up mechanisms proposed in (23). Our work not only provides a robust platform for exploring audio-tactile interactions but also sets the stage for investigating other cross-sensory integrations, enhancing our fundamental understanding of multisensory processing.

## Materials and methods

### Psychophysics Experiments

Six volunteers (3 male, mean age 33.83 *±* 6.74 SD) were recruited for Experiment 1 and twelve volunteers (7 male, mean age 35.5 *±* 10.72 SD) were recruited for Experiment 2 from the University of Exeter. Each gave written informed consent and received minor monetary compensation for participating in a 1-hour session. Participants were naive to the purpose of the study and did not self-declare any neurological or sensory disorders. Procedures were in compliance with guidelines for research with human participants and approved by the University of Exeter Research Ethics Committee.

Here we used the auditory streaming paradigm (49) in which participants are listening to a sequence of interleaved high and low-frequency tones repeated in ABA-triplets (“A” and “B” tones, “-” silent gap) (Fig 1A). They were instructed to report the “integrated” percept when they perceived these sequences either as a single integrated stream (A B A - A B A - A B A -), and the “segregated” percept when they heard two distinct streams: one consisting of only A tones and the other of only B tones (concurrent: A - A - A - A - A - A - and - B - - -B - - - B - -) (Fig 1B). Participants sat in a sound-isolated booth and attended to auditory stimuli while vibrotactile stimulators were attached to their left index finger. To ensure that participants fully understood these interpretations, auditory and visual demonstrations were provided. Participants were instructed to report their perceptions passively, without trying to favor one organization over the other. They used keyboard presses to indicate their perceptual responses.

**Figure 1:**
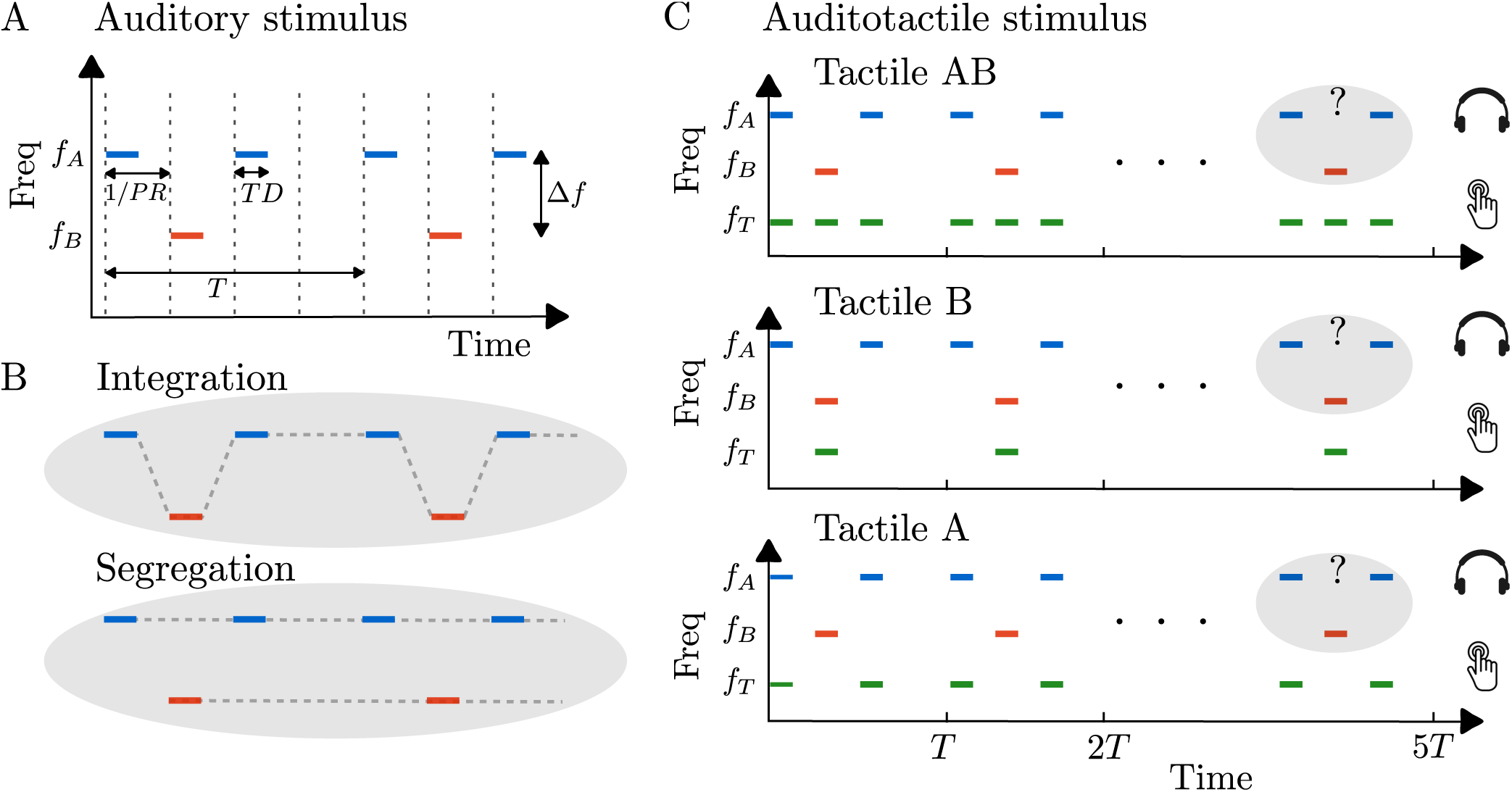
Auditory and tactile stimulus paradigm with two possible percepts. **A:** Repeating ABA-triplet sequences are composed of higher-frequency pure tones (A) interleaved with lower-frequency pure tones (B), each with a duration *TD*, and separated by a frequency difference Δ*f* . The time interval between successive tone onsets (indicated by dashed vertical lines) corresponds to the inverse of the presentation rate (1*/PR*). The “-” in ABA-represents a silent interval of duration 1*/PR*. In this study, tone duration is set to *TD* = 1*/PR*, ensuring that the offset of an A tone aligns precisely with the onset of the subsequent B tone. **B:** The stimulus is perceived in one of two ways: either as an integrated single stream (ABA-ABA-) or as two segregated streams (A-A-A-A- and -B—B–). Each experimental trial consists of five consecutive ABA-triplets, and participants were asked to report their perception as integrated or segregated for the last triplet of the 5-triplet sequence. **C:** In the Tact AB condition, tactile stimulations at *f_T_*occur during both A- and B-tone intervals. In contrast, in the Tact B and Tact A conditions, tactile stimulations occur only during B-tone or A-tone intervals, respectively.

The auditory stimuli consisted of five consecutive ABA-triplets, where each triplet included 125 *ms* pure tones (A and B) followed by a 125 *ms* silence (“-”), resulting in a total triplet duration of 500 *ms*. Cosine-squared ramps of 10 *ms* were applied to the onset and offset of each tone to ensure smooth transitions and avoid acoustic artefacts. The higher frequency A tones were a variable Δ*f* semitones (st) above the lower frequency B tones. A minimum 2 *s* interval between trials was used after which participants could run the next trial when ready.

Vibrotactile stimuli consisted of 125 *ms* sinusoidal vibratory pulses at 200 *Hz*. We used tactile stimulation synchronised with a subset of the tones in an ABA-triplet. Three different tactile pulse timings were considered (Fig 1C): one trial with tactile pulses synchronized with A tones (Tact A), one with tactile pulses synchronized with B tones (Tact B), and one with tactile pulses synchronized with both A and B tones (Tact AB). Additionally, a trial without tactile stimulation (audio only, Tact Off) was included. Participants were instructed to focus on auditory stimuli and report their perceptions based on the final auditory triplet presented in each trial.

The sequence of tones was played binaurally through Sennheiser HD 400 PRO headphones. We used miniature vibrotactile electromagnetic solenoid-type stimulators (18 *mm* diameter, Dancer Design tactors (50)) driven by a tactile amplifier (Dancer Design Tactamp (50)) to deliver tactile stimuli. In Experiment 1, each of the 6 participants completed 240 trials, consisting of 20 repetitions for each of three tactile conditions (Tact B, Tact AB, and Tact Off), combined with four different frequency difference levels (Δ*f* = *{*3, 4, 5, 6*} st*). We used a 6 × 6 Latin square design with 40 randomized and unique grids so that the order of conditions for each participant in a block of trials was counterbalanced within/across participants and block repetitions (one block of 6 participants and 40 repetitions). In Experiment 2, each of the 12 participants completed 160 trials, consisting of 40 repetitions for each of the four tactile conditions (Tact B, Tact AB, Tact Off, and Tact A) at a fixed frequency difference (Δ*f* = 4 *st*). Here we used a 4 × 4 Latin square design with 120 randomized and unique grids (three blocks of 4 participants and 40 repetitions).

### Statistical analysis

To investigate the effects of frequency difference (Δ*f*) and tactile condition on the proportion of trials reported as segregated, we employed a generalized linear mixed model (GLMM) with a logit link function (51), accounting for repeated measures within participants. This statistical analysis method is useful for modelling non-normally distributed responses, such as binary outcomes (segregated, not segregated), and incorporates both fixed effects and a random intercept for participants to account for individual variability (52). The fixed effects in the model estimate population-level coefficients, representing the relationship between predictor variables and the response, while random effects control for individual variations.

Different GLMMs were fitted to explore the main effects of Δ*f* , tactile conditions, and their interaction. The GLMM provides coefficient estimates that indicate the strength and direction of the effect of predictor variables on the outcome; a positive coefficient indicates that an increase in the predictor variable raises the likelihood of the outcome occurring, while a negative coefficient suggests the opposite. The absolute value of the coefficient represents the strength of the relationship, with larger values indicating a stronger effect. The significance level of 0.05 is used throughout this paper. All statistical analyses were conducted in the statistical package *R*. To estimate Cohen’s effect size (53), we first compute the odds ratio (OR) based on the observed frequencies of the binary outcome variable in Experiment 1. This odds ratio compares the odds of the event occurring in the test group to those in the control group. For the sample size estimation, we used G*Power (54) with a z-test for the difference between two independent proportions, employing an allocation ratio of 1 to achieve 80% power.

### Mathematical model for audio-tactile interaction

To investigate the effect of tactile pulses on the segregation of auditory streams, we developed a mechanistic mathematical model that captures interactions within auditory and tactile neural circuits (see Fig 3 for a schematic of the model). The neuronal circuits for competition and perceptual encoding included in the model are assumed to be downstream and receive inputs from the primary auditory cortex (A1). Neuronal activity is represented by mean firing rates, with competitive interactions arising through excitatory and inhibitory connections, slow adaptation, synaptic depression, and intrinsic noise.

The model is described by the following system of first-order differential equations based on the model presented in Rankin et al. (2015) (43), which is considered a discrete idealization of a tonotopically organized array:

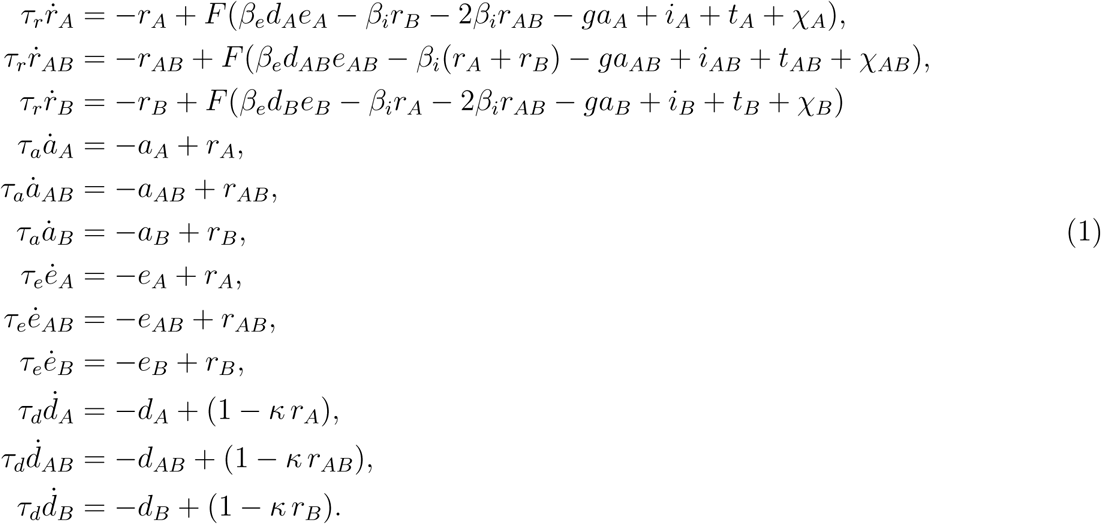

Here, *r_k_* are the mean firing rates of each population, indexed by *k* = *{A, AB, B}*. Dynamics of each population depends also on, spike frequency adaptation variable *a_k_*, recurrent NMDA-excitation variable *e_k_*, and synaptic depression of excitatory connections *d_k_*. *τ_r_*, *τ_a_* , *τ_e_* and *τ_d_* are synaptic time constants of respective variables. *β_e_* is the strength of recurrent NMDA-excitation *e_k_*, which is modulated by the slow synaptic depression *d_k_*. *β_i_* is the strength of instantaneous inhibition by the other populations *r_k_*. Inhibition from the *r_AB_* unit to the *r_A_* and *r_B_* units is twice as strong. *g* is the strength of spike-frequency adaptation, *a_k_*. *κ* is the strength of the slow synaptic depression.

Each population shares a sigmoidal firing rate function *F* , given by

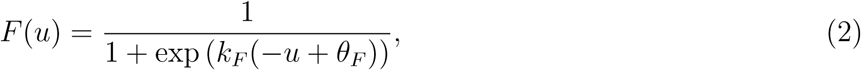

with threshold *θ_F_* and slope *k_F_* . The model is driven by periodic inputs that replicate the tonotopic responses in A1 to ABA-sequences (55). *i_k_* the tonotopic responses in the auditory cortex driving downstream neural populations are given by:

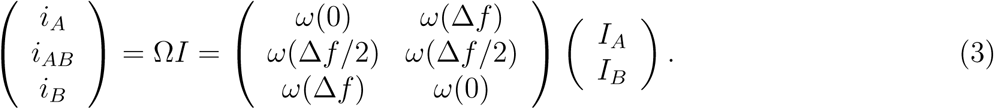

The spread of auditory *I_A_* and *I_B_* is defined by auditory weighting function 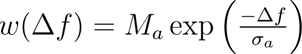, where *σ_a_* is a spatial decay parameter and *M_a_* is the pulse amplitude. The input terms *I_k_* (*k* = *{A, AB, B}*) mimic the onset-plateau responses to pure tones in A1 with onset timescale *α*_1_, plateau timescale *α*_2_ and peak to plateau ratio Λ_2_. Inputs are given by the following double *α*-function where *H*(*t*) is the Heaviside function:

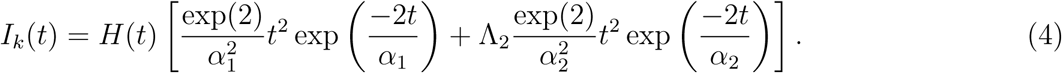

The interactive effect of tactile input on the downstream network is considered additive with inputs *t_k_* given by:

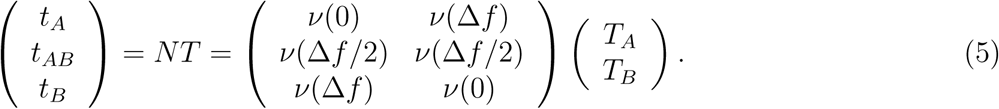

Input terms *T_A_*and *T_B_* represent tactile pulses that are synchronized with tone A and tone B, respectively. These inputs are defined by the double *α*-function given in Eq 4. The spread of these tactile pulses across tonotopic locations is defined by the tactile weighting function *ν*(Δ*f*) = 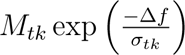, with parameters *σ_t_*_1_ and *M_t_*_1_ when tactile pulses are synchronized with one tone (Tact A or Tact B), and parameters *σ_t_*_2_ and *M_t_*_2_ when tactile pulses are synchronized with both A and B tones (Tact AB).

Additive noise is introduced with independent stochastic processes *χ_A_*, *χ_B_* and *χ_AB_* and added to the inputs of each population. Input noise is modelled as an Ornstein-Uhlenbeck process:

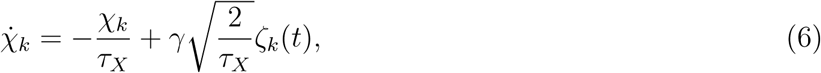

where *τ_X_*is the timescale, *γ* the strength and *ζ*(*t*) a white noise process with zero mean.

Values of all model parameters defined throughout this section are given in Table 1. Simulations were run in Matlab using a standard Euler-Murayama time stepping scheme with a stepsize of 1 ms. Reducing this stepsize by a factor of 10 did not change the results.

**Table 1:**
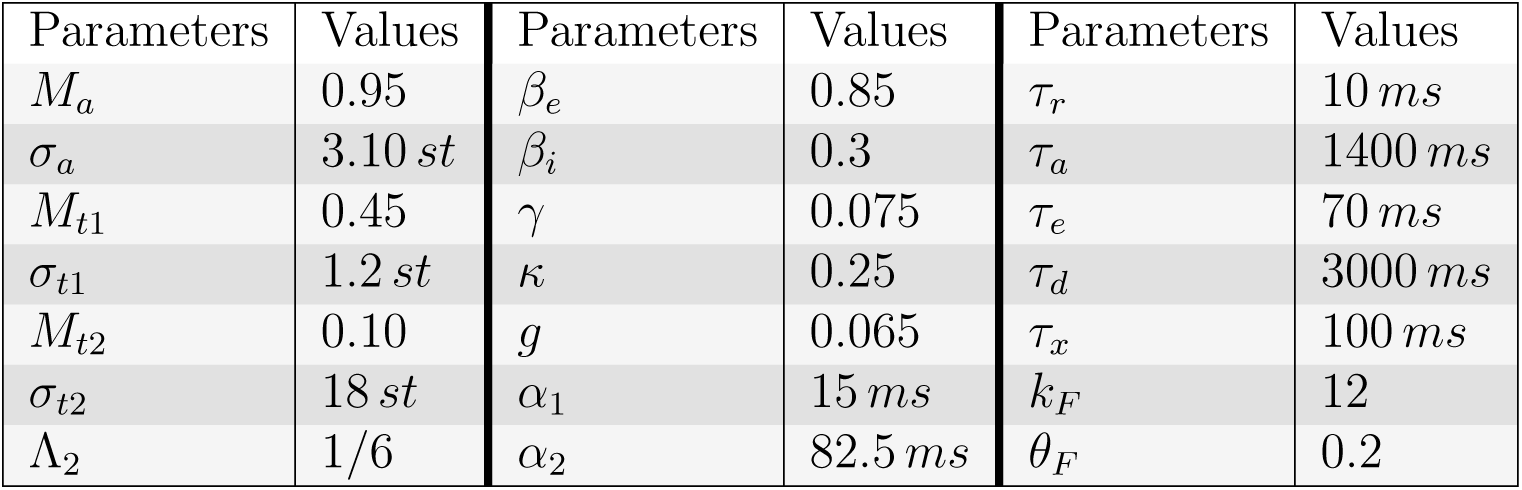
Parameters of the model with their corresponding values used in the simulations.

## Results

### Effect of tactile stimulation on perceptual segregation at different frequency differences

In Experiment 1 we investigate the effect of tactile pulses on the ability to segregate sound streams. In the experiment, participants listen to a sequence of five auditory triplets, consisting of interleaved high- and low-frequency tones arranged in an ABA-pattern (Fig 1A). They report whether they perceive the tones as a single, integrated stream or as segregated into two streams, based on their perception of the final triplet. We used tactile stimulation synchronised with a subset of the tones in an ABA-triplet (Fig 1C). Data were collected from 6 participants, repeating 20 times each of 12 conditions (1440 observations in total), with each participant reporting their perception across various conditions and frequency differences. When tactile stimulation timing matches only the B tone sequence, the proportion of trials in which participants report perceiving the sounds as segregated increases. When the tactile pulse timing matches the A and B tones a bias towards integration emerges. This effect was observed over a range of frequency difference values (Fig 2A).

**Figure 2:**
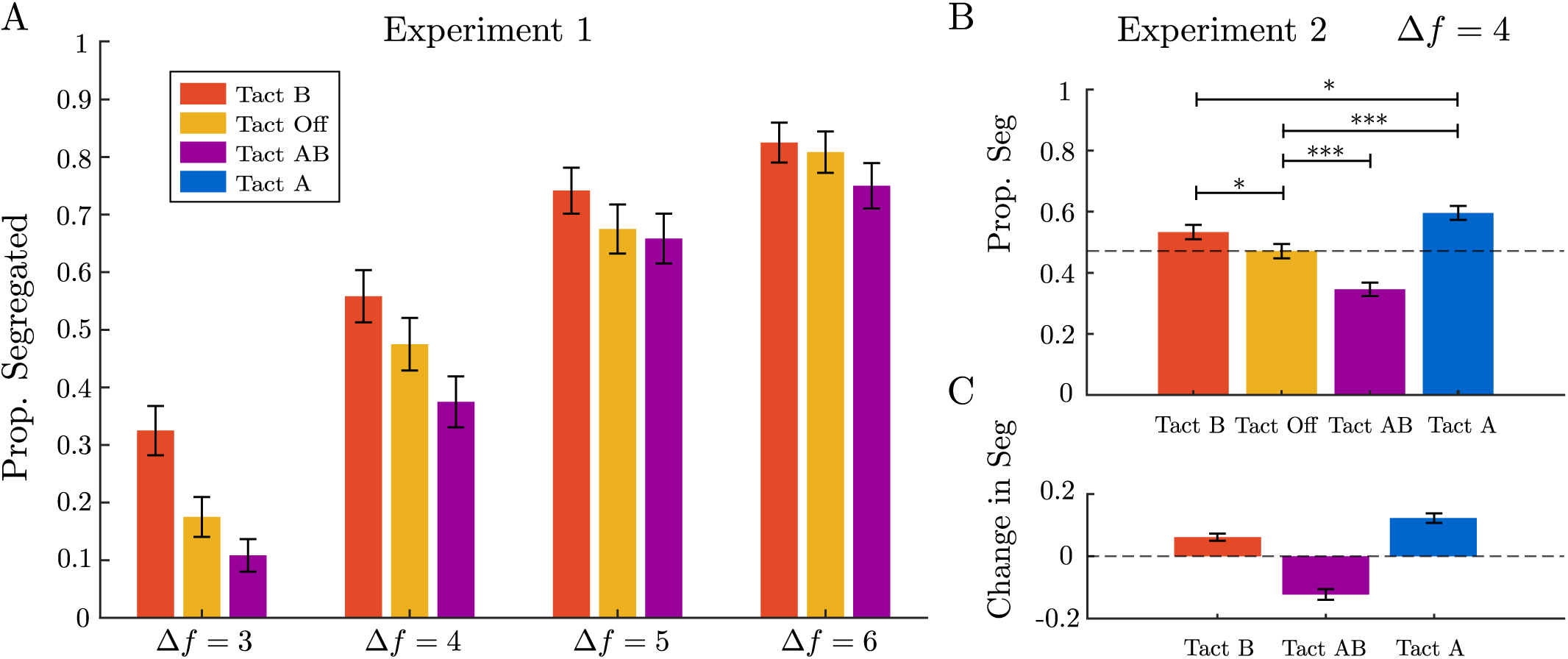
Effects of frequency difference, tactile conditions, and replicability. **A:** Experiment 1 (Combined effect of frequency difference and tactile conditions): Data were collected from six participants, each completing 20 repetitions at four frequency difference levels (Δ*f* = *{*3, 4, 5, 6*}*) crossed with three tactile conditions, Tact B, Tact Off, Tact AB. Participants reported their perception across various tactile conditions and frequency differences. Proportion segregated increases with Δ*f* and increases (decreases) in Tact B (Tact AB) relative to Tact Off. Error bars show the standard error of the proportion segregated. **B:** Experiment 2 (replicability, model validation): This experiment examines a fixed frequency difference (Δ*f* = 4) and includes an additional tactile condition in which tactile pulses are aligned with tone A in ABA triplets. Compared to the reference condition Tact Off, the condition Tact B had a marginally significant positive effect, while Tact A and Tact AB exhibited significant positive and negative effects, respectively. Participants’ perception of segregation was significantly lower in Tact B compared to Tact A. **C:** The change in proportion segregated compared to reference condition Tact Off.

To investigate the relationship between frequency difference (Δ*f*) and tactile condition and their effect on the perception of segregation in participants, we used a generalized linear mixed model (GLMM) analysis. The fixed effects of Δ*f* and tactile condition and their interaction were examined to assess how they influenced the likelihood of segregation. A random intercept for participants was included to account for individual variability. The analysis revealed significant fixed effects for both Δ*f* and tactile conditions. Specifically, the coefficient for Δ*f* (Coeff. estimate =1.01, *p <* 0.001) indicates a strong positive relationship between frequency difference and the likelihood of perceiving segregation; demonstrated in other studies without tactile stimulation (4; 3; 43). Additionally, Tact B showed a positive effect (Coeff. estimate=1.28, *p* = 0.047), while Tact AB exhibited a negative effect (Coeff. estimate = -0.64, *p* = 0.35). Notably, the interaction terms between Δ*f* and the tactile conditions did not reach statistical significance, suggesting that tactile conditions have a consistent effect and do not vary significantly across different frequency differences (Table 2). A simpler model without the interaction terms showed significant main effects for Δ*f* (Coeff. estimate=0.96, *p <* 0.001) and both tactile conditions Tact B (Coeff. estimate=0.42, *p* = 0.005) and Tact AB (Coeff. estimate=-0.32, *p* = 0.033); see Table 3.

**Table 2:** The coefficients of GLMM of the effects of frequency difference (Δ*f*) and tactile conditions on the segregation perception and their interactions while accounting for random effects of participants. Significant fixed effects are indicated with (*) or (***). The intercept and Δ*f* are showing highly significant effects. The interaction between Δ*f* and tactile conditions is not statistically significant. The AIC=1636.6 and BIC=1673.5 values suggest a good model fit.

| Variable | Estimate | Std. Error | z value | P-value |
| --- | --- | --- | --- | --- |
| (Intercept) | -4.36 | 0.50 | -8.72 | < .001 *** |
| $\Delta f$ | 1.01 | 0.10 | 9.70 | < .001 *** |
| Tact AB | -0.64 | 0.68 | -0.94 | 0.350 |
| Tact B | 1.28 | 0.64 | 1.99 | 0.047 * |
| $\Delta f$:Tact AB | 0.07 | 0.15 | 0.47 | 0.638 |
| $\Delta f$:Tact B | -0.20 | 0.14 | -1.39 | 0.166 |

**Table 3:** The coefficients of GLMM of the effects of frequency difference (Δ*f*) and tactile conditions on the segregation perception while accounting for random effects of participants. The model indicates a significant positive effect of Δ*f* and varying effects of the tactile conditions, with Tact AB showing a significant negative estimate and Tact B indicating a significant positive effect. Lower Std. Error and slightly lower BIC=1662.6 value suggest a better model fit compared to the model including interactions (AIC=1636.3).

| Variable | Estimate | Std. Error | z value | P-value |
| --- | --- | --- | --- | --- |
| (Intercept) | -4.16 | 0.33 | -12.66 | < .001 *** |
| $\Delta f$ | 0.96 | 0.06 | 16.05 | < .001 *** |
| Tact AB | -0.32 | 0.15 | -2.13 | 0.033 * |
| Tact B | 0.42 | 0.15 | 2.81 | 0.005 ** |

Analyzing the data at a specific frequency difference (Δ*f* = 4) showed that Tact B had a positive effect on segregation perception compared to Tact Off (Coeff. estimate=0.36, *p* = 0.179), whereas Tact AB exhibited a negative effect (Coeff. estimate = -0.45, *p* = 0.103). However, neither effect reached statistical significance (Table 4). The odds ratios of Tact AB and Tact B compared to the reference level of Tact Off are equal to 0.64 and 1.44, corresponding to Cohen’s d (log odds ratio) 0.25 and 0.20 respectively. We used the smaller effect size of 0.2 of the tactile stimulation in Experiment 1 to estimate the sample size of a follow-up experiment (Experiment 2). To detect an effect size of 0.2 with 80% power using a z-test for the difference between two independent proportions, G*Power (54) estimates a required sample size of at least 444 (equivalent to 12 participants completing 40 repetitions for each condition).

**Table 4:** The coefficients of GLMM of the influence of tactile conditions on perceptual segregation at a fixed frequency difference (Δ*f* = 4) in Experiment 1. Fixed effects Tact AB, and Tact B reveal no statistically significant differences in perceptual segregation between the conditions

| Variable | Estimate | Std. Error | z value | P-value |
| --- | --- | --- | --- | --- |
| (Intercept) | -0.10 | 0.32 | -0.31 | 0.757 |
| Tact AB | -0.45 | 0.27 | -1.63 | 0.103 |
| Tact B | 0.36 | 0.27 | 1.34 | 0.179 |

Results from Experiment 1 reveal that tactile pulses have an effect on the ability to segregate auditory streams. Tactile pulses synchronized with tone B promote segregation, while those synchronized with both A and B tones promote integration. This suggests a complex interaction between tactile and auditory modalities, where the timing and context of tactile input are crucial. In Experiment 2 we investigate tactile stimulation at a fixed frequency difference (Δ*f* = 4, close to equidominance point) to better understand the effects of temporal alignment of tactile and auditory stimuli. To this end, Experiment 2 also includes an additional tactile condition in which tactile pulses are aligned with A tones in ABA-triplets (in this way participants are exposed to twice as many tactile stimuli as in Tact B condition). The results of the experiment are shown in Fig 2B. The change in the proportion segregated for each tactile condition compared to the reference condition Tact Off is depicted in Fig 2C. Values greater than zero indicate a bias towards segregation, while values less than zero indicate a bias towards integration.

Compared to the reference condition Tact Off, Tact AB had a significant negative effect (Coeff. estimate = -0.54, *p <* 0.001), while Tact A (Coeff. estimate = 0.53, *p <* 0.001) and Tact B (Coeff. estimate = 0.26, *p* = 0.048) exhibited significant positive effect; see also Table 5. Compared to Tact B, Tact A demonstrates a significantly higher proportion of segregation (Coeff. estimate = 0.27, *p* = 0.046).

**Table 5:** The coefficients of GLMM of the influence of tactile conditions on perceptual segregation at a fixed frequency difference (Δ*f* = 4) in Experiment 2.

| Variable | Estimate | Std. Error | z value | P-value |
| --- | --- | --- | --- | --- |
| Tact AB - Tact Off | -0.54 | 0.14 | -4.00 | < .001 *** |
| Tact B - Tact Off | 0.26 | 0.13 | 1.98 | 0.048 * |
| Tact A - Tact Off | 0.53 | 0.13 | 3.95 | < .001 *** |
| Tact A - Tact B | 0.27 | 0.13 | -2.00 | 0.046 * |
| Tact B - Tact AB | 0.80 | 0.14 | 5.92 | < .001 *** |
| Tact A - Tact AB | 1.07 | 0.14 | 7.81 | < .001 *** |

### Mechanistic mathematical model of tactile-induced bias in auditory stream segregation

The model includes three neuronal populations. Two of them pool inputs from A1 regions centred on the frequencies A and B. The third population receives input from an intermediate tonotopic location, approximately (*A* + *B*)*/*2; see Fig 3A). The specific frequency tones exhibit full amplitude at their respective tonotopic locations, gradually decaying as they spread across other tonotopic locations (decay function *ω*(Δ*f*)). Similarly, the effect of tactile pulses is assumed to diminish across tonotopic locations, as illustrated by the changes in inputs corresponding to Tact B (represented by dashed curves in Fig 3B). When tactile pulses are synchronized with B-tones, they enhance the tonotopic response at location B and also induce an excitatory effect on other locations (AB and A), though to a lesser extent. We assumed a similar pattern of excitation for tactile pulses synchronized with A tones (decay function *ν*(Δ*f*)). However, when tactile pulses were synchronized with both tones, we considered a decaying effect, modelled with a different profile, to account for any residual impact from one tactile pulse to the next, as they occur at a shorter temporal distance. The top panel in Fig 3C shows a model simulation of the recurrent excitation variables for each population without the tactile effect (Tact Off). When the central AB unit is active (integrated), peripheral units are suppressed through mutual inhibition. Increasing adaptation for AB raises the probability of noise-induced switching, leading to the activation and dominance of units A or B (segregated), which in turn suppresses the integrated (AB) unit. The panels below depict predicted percepts under various tactile conditions, showing that Tact B increases the segregation proportion, while Tact AB decreases it compared to the Tact Off condition.

**Figure 3:**
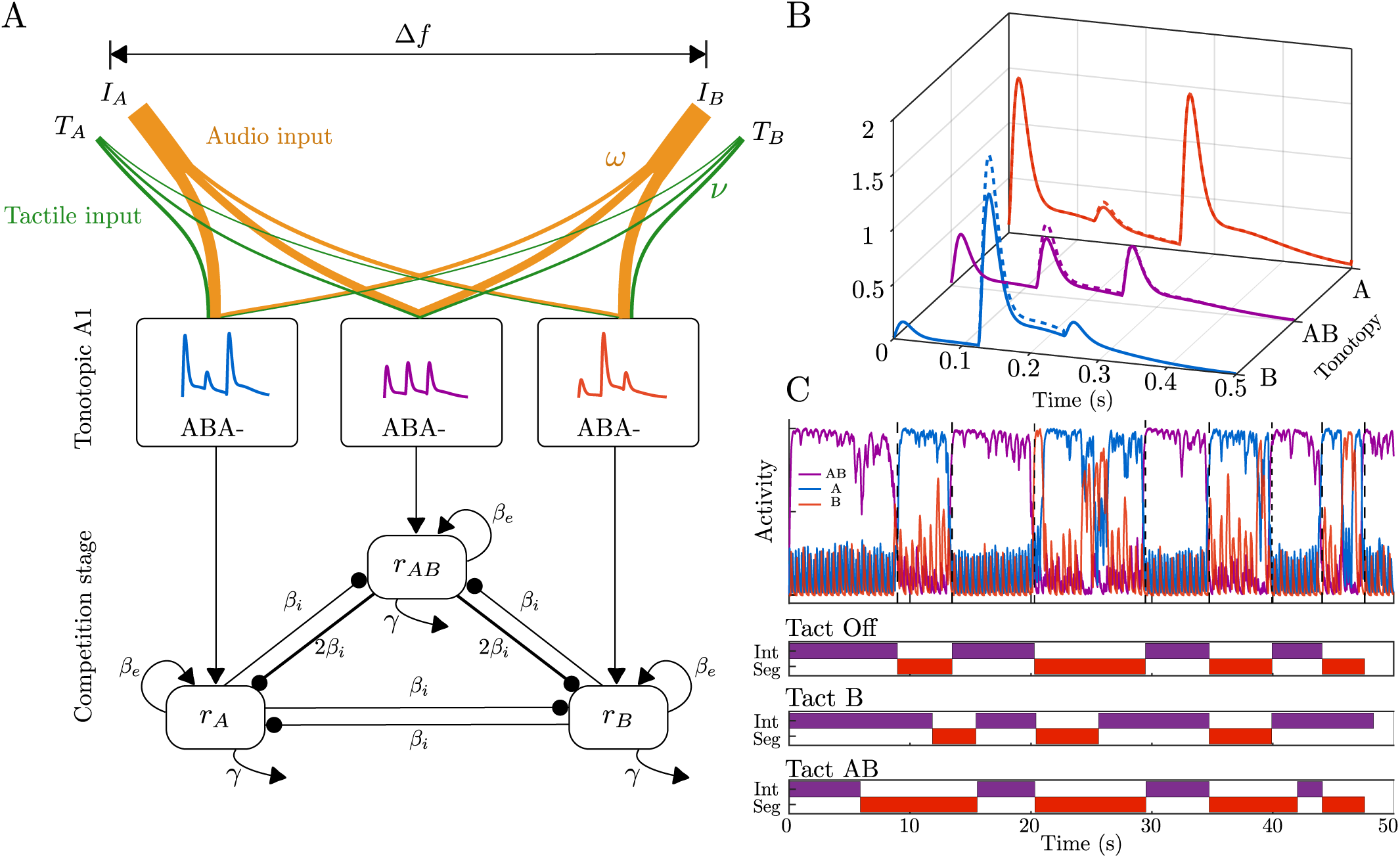
Model architecture, inputs, time course of model responses and predicted percepts. **A:** Neuromechanistic model with competition between units (*r_A_*, *r_AB_*, *r_B_*) driven by inputs from three locations in tonotopic map in A1 (at A, B and a location in between). The competition stage is located downstream of (and takes input from) A1, with mutual inhibition between units (*β_i_*), recurrent excitation (*β_e_*), slow adaptation (*γ*) and noise-driven competition. **B:** Inputs to the respective populations *r_A_*, *r_AB_* and *r_B_* for an ABA-triplet which shows the spread of inputs across the model’s tonotopy. The A-tone and B-tone inputs (*I_A_* and *I_B_*) have full amplitude at their respective tonotopic locations and gradually decay as they spread across other tonotopic locations (solid curves). The effect of tactile pulses is also considered to decay across tonotopic locations (changes in inputs corresponding to Tact B represented by dashed curves). **C:** The top panel displays a model simulation of the recurrent excitation variables for each population without tactile effect (Tact Off). When the central AB unit is active (purple/ integrated), peripheral units are suppressed via mutual inhibition. Increasing adaptation in the AB unit raises the likelihood of noise-induced switching. The panels below show predicted percepts for different tactile conditions with tact B (Tact AB) increasing (decreasing) the segregation proportion compared to Tact Off condition.

To investigate the mechanism of the effect of tactile pulses on the segregation of auditory streams we checked if the model presented in the section “Mathematical model for audio-tactile interaction” can be used to reproduce experimental observations. To this end, we used a genetic algorithm optimization implemented in Matlab function ga to find parameters *M_a_, M_t_*_1_*, M_t_*_2_ and *σ_a_, σ_t_*_1_*, σ_t_*_2_ of the *ω*(Δ*f*) and *ν*(Δ*f*) functions; used in Eqs (3 and 4), respectively. The other model parameters were fixed throughout the optimization procedure. We defined the cost function as the mean squared error between the proportions of segregation observed in experiments and the proportions of segregation obtained in simulations. Details of the optimization procedure can be found in the computer code associated with the paper.

Results of the simulations for the optimal parameter values are shown in Fig 4. Experimental data are presented as bar plots, while results of the model simulations are shown as dashed curves with data points at different frequency differences (Δ*f* = *{*3, 4, 5, 6*} st*) and varying tactile conditions (Fig 4A). The Δ*f* -dependent profiles for the auditory input spread *ω*(Δ*f*) (Fig 4C) and tactile input spread *ν*(Δ*f*) (Fig 4D) were determined through the optimization algorithm. The decaying input function was estimated to have an amplitude of *M_a_* = 0.95 *±* 0.04 and a slope of *σ_a_* = 3.10 *±* 0.12. For the case where tactile pulses are aligned with one tone, the parameters are *M_t_*_1_ = 0.45 *±* 0.12 and *σ_t_*_1_ = 1.2 *±* 0.22. When tactile pulses are aligned with both tones, the amplitude is *M_t_*_2_ = 0.10 *±* 0.04 and the slope is *σ_t_*_2_ = 18*±*7. Confidence intervals are based on 20 repetitions of the fitting procedure. The effect of tactile input amplitude is smaller compared to auditory input, and the decay across the tonotopic representation is steeper in single-tone alignment than in auditory decay, while remaining almost uniform in dual-tone alignment.

**Figure 4:**
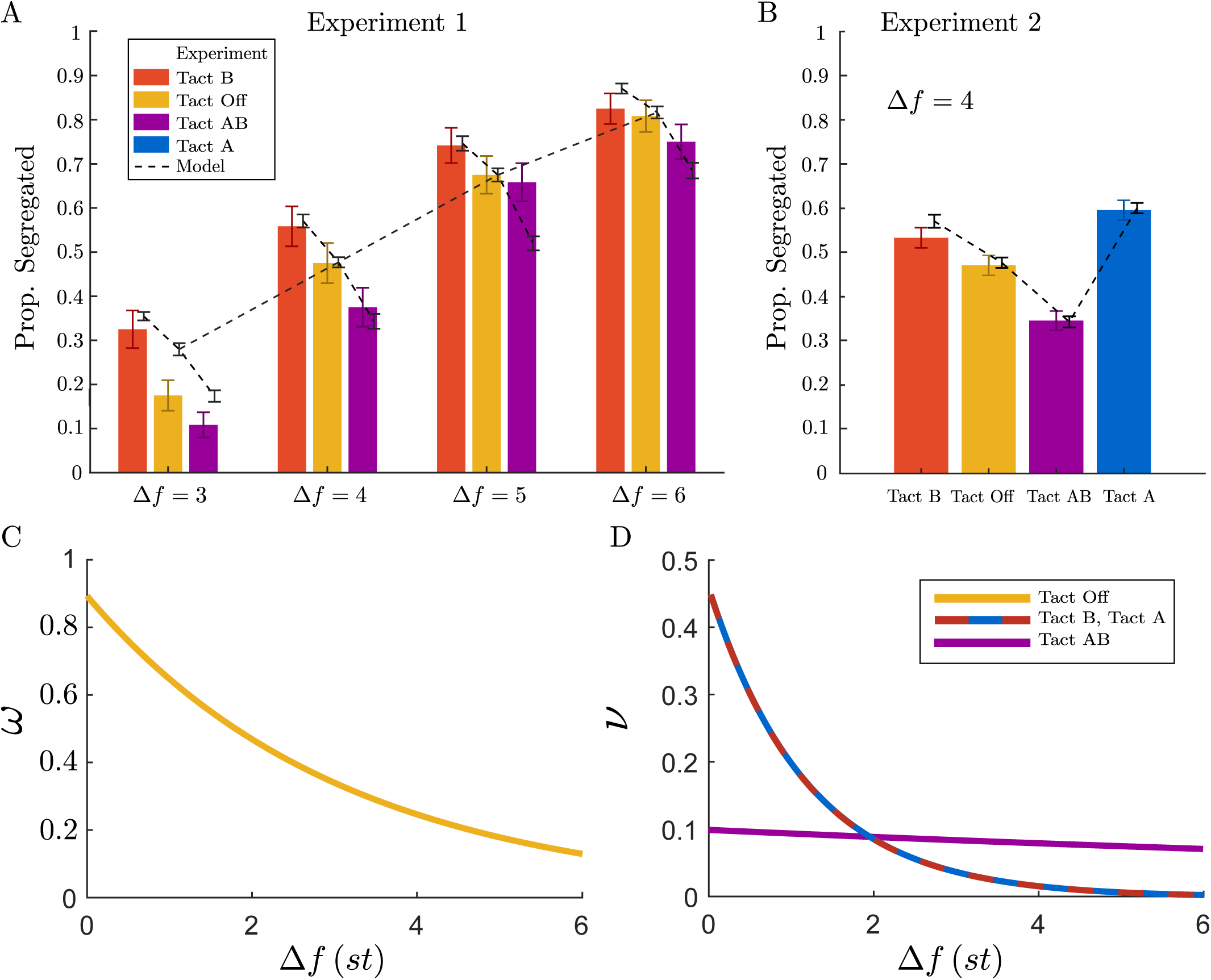
Comparison of experimental and simulated data across frequency differences and tactile Conditions, with audio and tactile input spread profiles. **A-B:** Experimental data are presented as bar plots, while simulated data are shown as dashed curves with data points at different frequency differences (Δ*f* = 3, 4, 5, 6 *st*) and under different tactile conditions. Error bars show the standard error of the proportion segregated. **C-D:** The Δ*f* -dependent profiles for the auditory input spread *ω*(Δ*f*) and tactile input spread *ν*(Δ*f*) (modelled as exponential decays) were derived using an optimization algorithm minimizing the mean squared error between experimental and computational data from Experiment 1. These weighting functions were then used to compare computational model predictions with experimental data in Experiment 2.

We used model parameters estimated using data from Experiment 1 to run a new set of simulations that included tactile stimulus synchronized with tone A. We compared the results of the new simulation with data from Experiment 2. Without any additional fitting, the model made an accurate quantitative prediction that the level of segregation under the new tactile condition, Tact A, is higher than Tact B. Fig 4B illustrates excellent alignment between the model and the experimental data.

## Discussion

### Tactile pulses interfere with tonotopic responses

By synchronizing tactile pulses with specific tones in the ABA-triplet sequence, we explored the perceptual shifts between integration (one stream) and segregation (two streams) based on the timing and pattern of tactile stimulation. Our findings imply that cross-modal synchronization can dynamically shift perceptual boundaries in auditory streaming. When tactile pulses coincided only with tone B, experiments demonstrated an increased likelihood of segregating the tones. When both A and B tones were paired with tactile pulses, participants perceived the sounds as a single stream more often. This suggests that tactile stimulation synchronized with a single tone in the sequence enhances perceptual distinction between tones, reinforcing the auditory separation needed for segregation, while alignment of both tones and tactile pulses provides a unified cross-modal cue, leading the auditory system to interpret the sequence as more cohesive. Tactile conditions have a consistent effect across various frequency differences suggesting that tactile timing relative to sound structure is a strong factor in stream segregation.

### Tonotopic modulation and temporal averaging

In our model, tactile pulses modulate auditory perception by altering tonotopic responses in the auditory cortex. In the primary auditory cortex, neurons that respond to similar frequencies are organized in clusters along the tonotopic map (56). Excitatory neurons activate neighbouring frequency-tuned neurons, while inhibitory interneurons create lateral inhibition, sharpening frequency specificity and limiting spread across the map (57). When tactile pulses align with a single tone, they appear to selectively boost responses in regions associated with that tone, with the effect decaying across the tonotopic map. This supports segregation by enhancing contrast in tonotopic representation. When both tones are paired with tactile stimuli, broader, more uniform activation across tonotopic areas reduces this contrast, promoting integration.

The increased segregation observed when tactile pulses align with tones A in the ABA-triplets, compared to B tones, is particularly notable due to the presence of two tactile pulses instead of one. With two tactile stimuli per ABA triplet, the tactile impact curve decays as sharply as before. However, because the tactile stimuli are now repeated twice within a triplet, they reinforce the auditory processing of tone A, leading to a more pronounced segregation effect.

We hypothesize that the observed effect could be explained through neural mechanisms in the auditory cortex. When tactile pulses synchronize with auditory tones, they interact with tonotopic circuits, activating frequency-tuned neurons and spreading in a balanced manner controlled by excitation and inhibition. When tactile pulses activate neurons associated with both tones, the individual effects integrated in time lead to a combined activation in the neural responses that do not decay as sharply across the tonotopic map. This could be due to recurrent excitation, where the remaining activity from one pulse keeps the neural population primed, leading to a more sustained response. The combination of effects at both tonotopic locations could create a stable perceptual outcome that enhances integration, as the stimulation is spread out more evenly (58).

Alternatively, the experimental observations could be explained by temporal averaging of excitatory and inhibitory effects. Such averaging would lead to the tactile impact curve that is smaller and smoother when tactile pulses align with both tones versus just one. When tactile pulses are synchronized with both tones, the excitatory and inhibitory responses may partially overlap and combine across time, leading to a more balanced, averaged response across the tonotopic map. This smoothing effect would reduce the peak strength of activation but maintain it across a wider spatial area, resulting in a smaller yet more stable effect. In contrast, with tactile pulses aligned with one tone, the activation is enhanced for that specific frequency region. Such localized amplification could explain the stronger but spatially limited effect in the one-tone condition.

### Addtive tactile effect and pattern of tactile pulses

Using the statistical GLM model we observed no significant interaction between frequency difference (Δ*f*) and tactile pulses. The lack of significant interaction terms between Δ*f* and tactile conditions suggests that tactile inputs have a consistent influence that does not vary notably with different frequency differences. These findings justify, modelling the tactile stimulations as purely additive effects. This approach provides a simplified framework to capture the subtle tactile effects without introducing complex interactions.

### Future work and applications

Previous studies have demonstrated that tactile distractors can bias auditory frequency perception, shifting perceived frequencies toward those of the tactile pulses (20). This bias is more pronounced with lower-frequency tactile distractors compared to higher frequencies. In this study, we specifically used tactile pulses with frequencies lower than the auditory tones. Future research could investigate the impact of higher-frequency tactile pulses on auditory perception to determine if they produce a different bias or interaction pattern, deepening our understanding of multisensory integration dynamics across varying frequency contexts.

Our experimental and mathematical modelling results reveal a spatio-temporal interaction where tactile input dynamically influences auditory organization through timing and tonotopic alignment. Specifically, our findings show that dual-tone alignment creates a broad, stable neural response, while single-tone alignment produces a sharper, more localized effect. This interaction suggests that tactile cues, precisely timed and placed, can be employed to fine-tune auditory processing. Such intricate control offers valuable insights into multisensory integration, especially for the design of devices for individuals with hearing impairments or difficulties with speech recognition in noisy environments. By strategically incorporating tactile feedback, our approach can improve and optimize the design of these devices, enhancing auditory clarity and speech recognition.

## Conclusion

This study highlights the significant role of tactile pulses in modulating auditory perception, demonstrating that synchronized tactile input can influence the integration and segregation of auditory streams through spatiotemporal interactions. These effects arise from interactions within tonotopic circuits, where excitatory and inhibitory dynamics shape the perceptual outcome. Single-tone alignment creates localized enhancement and dual-tone alignment generates broader, averaged responses. The consistency of tactile effects across varying frequency differences highlights the robustness of tactile-auditory interactions. These insights open pathways for future research to explore how tactile inputs of different frequencies influence auditory perception, further advancing our understanding of multisensory integration. Importantly, the ability to fine-tune auditory processing using precisely timed tactile cues has promising applications, particularly in assistive technologies. This work lays the foundation for innovative solutions that capitalize on multisensory integration to optimize auditory experiences.

## Acknowledgments

We would like to thank Marc Goodfellow and Mark Fletcher for their valuable discussions and insightful suggestions during the early stages of this project.

## Availability of data and materials

The source code for the model and the datasets generated and analyzed during the current study will be available in the GitHub repository at the time of publication.

## Contributions

All authors contributed to the experimental design, model development, theoretical analysis, and discussion of the results. FD implemented the model, conducted the experiments, analyzed the data, and wrote the manuscript. All authors reviewed and approved the final version of the manuscript.

## Ethics approval

Approval was obtained from the ethics committee of the University of Exeter (eEMPS000058). The procedures used in this study adhere to the tenets of the Declaration of Helsinki. Written formal consent was obtained from all individual participants included in the study.

## References

[1] Chung K. Challenges and recent developments in hearing aids: Part I. Speech understanding in noise, microphone technologies and noise reduction algorithms. Trends in Amplification. 2004;8(3):83–124.

[2] Snyder JS, Alain C. Toward a neurophysiological theory of auditory stream segregation. Psychological bulletin. 2007;133(5):780.

[3] Moore BC, Gockel HE. Properties of auditory stream formation. Philosophical Transactions of the Royal Society B: Biological Sciences. 2012;367(1591):919–31.

[4] van Noorden LPAS. Temporal coherence in the perception of tone sequences. 1975.

[5] Fishman YI, Reser DH, Arezzo JC, Steinschneider M. Neural correlates of auditory stream segregation in primary auditory cortex of the awake monkey. Hearing research. 2001;151(1-2):167–87.

[6] Carlyon RP, Plack CJ, Fantini DA, Cusack R. Cross-modal and non-sensory influences on auditory streaming. Perception. 2003;32(11):1393–402.

[7] Marozeau J, Innes-Brown H, Grayden DB, Burkitt AN, Blamey PJ. The effect of visual cues on auditory stream segregation in musicians and non-musicians. PLoS One. 2010;5(6):e11297.

[8] Slater KD, Marozeau J. The effect of tactile cues on auditory stream segregation ability of musicians and nonmusicians. Psychomusicology: Music, Mind, and Brain. 2016;26(2):162.

[9] Schulte A, Marozeau J, Ruhe A, Büchner A, Kral A, Innes-Brown H. Improved speech intelligibility in the presence of congruent vibrotactile speech input. Scientific Reports. 2023;13(1):22657.

[10] Fletcher MD, Verschuur CA, Perry SW. Improving speech perception for hearing-impaired listeners using audio-to-tactile sensory substitution with multiple frequency channels. Scientific Reports. 2023;13(1):13336.

[11] Fletcher MD, Akis E, Verschuur CA, Perry SW. Improved tactile speech perception and noise robustness using audio-to-tactile sensory substitution with amplitude envelope expansion. Scientific Reports. 2024;14(1):15029.

[12] Auvray M, Myin E. Perception with compensatory devices: From sensory substitution to sensorimotor extension. Cognitive Science. 2009;33(6):1036–58.

[13] Auvray M, Harris LR. The state of the art of sensory substitution. Multisensory research. 2014;27(5-6):265–9.

[14] Schwartz JL, Grimault N, Hupé JM, Moore BC, Pressnitzer D. Multistability in perception: binding sensory modalities, an overview. Philosophical Transactions of the Royal Society B: Biological Sciences. 2012;367(1591):896–905.

[15] MacDonald J, McGurk H. Visual influences on speech perception processes. Perception & psychophysics. 1978;24(3):253–7.

[16] King AJ. Visual influences on auditory spatial learning. Philosophical Transactions of the Royal Society B: Biological Sciences. 2009;364(1515):331–9.

[17] Opoku-Baah C, Schoenhaut AM, Vassall SG, Tovar DA, Ramachandran R, Wallace MT. Visual influences on auditory behavioral, neural, and perceptual processes: a review. Journal of the Association for Research in Otolaryngology. 2021;22(4):365–86.

[18] Bolognini N, Papagno C, Moroni D, Maravita A. Tactile temporal processing in the auditory cortex. Journal of Cognitive Neuroscience. 2010;22(6):1201–11.

[19] Hoefer M, Tyll S, Kanowski M, Brosch M, Schoenfeld MA, Heinze HJ, et al. Tactile stimulation and hemispheric asymmetries modulate auditory perception and neural responses in primary auditory cortex. Neuroimage. 2013;79:371–82.

[20] Yau JM, Weber AI, Bensmaia SJ. Separate mechanisms for audio-tactile pitch and loudness interactions. Frontiers in psychology. 2010;1:160.

[21] Cappe C, Rouiller EM, Barone P. Multisensory anatomical pathways. Hearing research. 2009;258(1-2):28–36.

[22] Klemen J, Chambers CD. Current perspectives and methods in studying neural mechanisms of multisensory interactions. Neuroscience & Biobehavioral Reviews. 2012;36(1):111–33.

[23] Kayser C, Petkov CI, Augath M, Logothetis NK. Integration of touch and sound in auditory cortex. Neuron. 2005;48(2):373–84.

[24] Foxe JJ, Schroeder CE. The case for feedforward multisensory convergence during early cortical processing. Neuroreport. 2005;16(5):419–23.

[25] Macaluso E, Driver J. Multisensory spatial interactions: a window onto functional integration in the human brain. Trends in neurosciences. 2005;28(5):264–71.

[26] Schroeder CE, Foxe J. Multisensory contributions to low-level,â€unisensoryâ€™processing. Current opinion in neurobiology. 2005;15(4):454–8.

[27] Raij T, Ahveninen J, Lin FH, Witzel T, Jääskeläinen IP, Letham B, et al. Onset timing of cross-sensory activations and multisensory interactions in auditory and visual sensory cortices. European Journal of Neuroscience. 2010;31(10):1772–82.

[28] Fu KMG, Johnston TA, Shah AS, Arnold L, Smiley J, Hackett TA, et al. Auditory cortical neurons respond to somatosensory stimulation. Journal of Neuroscience. 2003;23(20):7510–5.

[29] Schroeder CE, Lindsley RW, Specht C, Marcovici A, Smiley JF, Javitt DC. Somatosensory input to auditory association cortex in the macaque monkey. Journal of neurophysiology. 2001;85(3):1322–7.

[30] Schroeder CE, Smiley J, Fu KG, McGinnis T, O’Connell MN, Hackett TA. Anatomical mechanisms and functional implications of multisensory convergence in early cortical processing. international Journal of Psychophysiology. 2003;50(1-2):5–17.

[31] Foxe JJ, Wylie GR, Martinez A, Schroeder CE, Javitt DC, Guilfoyle D, et al. Auditory-somatosensory multisensory processing in auditory association cortex: an fMRI study. Journal of neurophysiology. 2002;88(1):540–3.

[32] Schürmann M, Caetano G, Hlushchuk Y, Jousmäki V, Hari R. Touch activates human auditory cortex. Neuroimage. 2006;30(4):1325–31.

[33] Yau JM, Olenczak JB, Dammann JF, Bensmaia SJ. Temporal frequency channels are linked across audition and touch. Current biology. 2009;19(7):561–6.

[34] Henschke JU, Noesselt T, Scheich H, Budinger E. Possible anatomical pathways for short-latency multisensory integration processes in primary sensory cortices. Brain Structure and Function. 2015;220:955–77.

[35] Fu X, Riecke L. Effects of continuous tactile stimulation on auditory-evoked cortical responses depend on the audio-tactile phase. NeuroImage. 2023;274:120140.

[36] Ranjbar P, Wilson EC, Reed CM, Braida LD. Auditory-tactile integration: Effects of phase of sinusoidal stimulation at 50 and 250 hz. International journal of engineering technology and scientific innovation. 2016;1(2):209.

[37] Lunghi C, Morrone MC, Alais D. Auditory and tactile signals combine to influence vision during binocular rivalry. Journal of Neuroscience. 2014;34(3):784–92.

[38] Jagt M, Ganis F, Serafin S. Enhanced neural phase locking through audio-tactile stimulation. Frontiers in Neuroscience. 2024;18:1425398.

[39] Szabó BT, Denham SL, Winkler I. Computational models of auditory scene analysis: a review. Frontiers in Neuroscience. 2016;10:524.

[40] Rankin J, Rinzel J. Computational models of auditory perception from feature extraction to stream segregation and behavior. Current opinion in neurobiology. 2019;58:46–53.

[41] Darki F, Ferrario A, Rankin J. Hierarchical processing underpins competition in tactile perceptual bistability. Journal of Computational Neuroscience. 2023;51(3):343–60.

[42] Snyder JS, Elhilali M. Recent advances in exploring the neural underpinnings of auditory scene perception. Annals of the New York Academy of Sciences. 2017;1396(1):39–55.

[43] Rankin J, Sussman E, Rinzel J. Neuromechanistic model of auditory bistability. PLoS computational biology. 2015;11(11):e1004555.

[44] Rankin J, Osborn Popp PJ, Rinzel J. Stimulus pauses and perturbations differentially delay or promote the segregation of auditory objects: psychoacoustics and modeling. Frontiers in Neuroscience. 2017;11:198.

[45] Ferrario A, Rankin J. Auditory streaming emerges from fast excitation and slow delayed inhibition. The Journal of Mathematical Neuroscience. 2021;11(1):8.

[46] Moreno-Bote R, Rinzel J, Rubin N. Noise-induced alternations in an attractor network model of perceptual bistability. Journal of neurophysiology. 2007;98(3):1125–39.

[47] Shpiro A, Moreno-Bote R, Rubin N, Rinzel J. Balance between noise and adaptation in competition models of perceptual bistability. Journal of computational neuroscience. 2009;27:37–54.

[48] Kondo HM, Pressnitzer D, Shimada Y, Kochiyama T, Kashino M. Inhibition-excitation balance in the parietal cortex modulates volitional control for auditory and visual multistability. Scientific reports. 2018;8(1):14548.

[49] Bregman AS. Auditory streaming: Competition among alternative organizations. Perception & Psychophysics. 1978;23:391–8.

[50] Dancer C. Dancer design company;. https://www.dancerdesign.co.uk/products.html.

[51] Parzen M, Ghosh S, Lipsitz S, Sinha D, Fitzmaurice GM, Mallick BK, et al. A generalized linear mixed model for longitudinal binary data with a marginal logit link function. The annals of applied statistics. 2011;5(1):449.

[52] Siegel LK, Silva M, Lin L, Chen Y, Liu YL, Chu H. Choice of link functions for generalized linear mixed models in meta-analyses of proportions. Research Methods in Medicine & Health Sciences. 2023:26320843231224808.

[53] Goulet-Pelletier JC, Cousineau D. A review of effect sizes and their confidence intervals, Part I: The Cohenâ€™sd family. The Quantitative Methods for Psychology. 2018;14(4):242–65.

[54] Faul F, Erdfelder E, Buchner A, Lang AG. Statistical power analyses using G* Power 3.1: Tests for correlation and regression analyses. Behavior research methods. 2009;41(4):1149–60.

[55] Micheyl C, Tian B, Carlyon RP, Rauschecker JP. Perceptual organization of tone sequences in the auditory cortex of awake macaques. Neuron. 2005;48(1):139–48.

[56] Langers DR, van Dijk P. Mapping the tonotopic organization in human auditory cortex with minimally salient acoustic stimulation. Cerebral cortex. 2012;22(9):2024–38.

[57] Wu GK, Arbuckle R, Liu Bh, Tao HW, Zhang LI. Lateral sharpening of cortical frequency tuning by approximately balanced inhibition. Neuron. 2008;58(1):132–43.

[58] Fishman YI, Kim M, Steinschneider M. A crucial test of the population separation model of auditory stream segregation in macaque primary auditory cortex. Journal of Neuroscience. 2017;37(44):10645–55.

